# ASM-3D: An attentional search model fashioned after what and where/how pathways for target search in 3D environment

**DOI:** 10.1101/2022.08.01.502278

**Authors:** Sweta Kumari, V Y Shobha Amala, M Nivethithan, V. Srinivasa Chakravarthy

**Affiliations:** PhD Scholar, Computational Neuroscience (CNS) Lab, Department of Biotechnology, IIT Madras, 600036, India; Intern, IIT BHU, 221005, India; Dual Degree, CNS Lab, Department of Biotechnology, IIT Madras, 600036, India; Professor, CNS Lab, Department of Biotechnology, IIT Madras, 600036, India

**Keywords:** Attention, Memory, Human Visual System, What and Where pathway, Convolutional neural network, Search in 3D, Flip-flop

## Abstract

We propose a biologically inspired attentional search model for target search in a 3D environment, which has two separate channels for object classification, analogous to the “what” pathway in the human visual system, and for prediction of the next location of the camera, analogous to the “where” pathway. We generated 3D Cluttered Cube datasets that consist of an image on one vertical face, and clutter images on the other faces. The camera goes around each cube on a circular orbit centered on the cube and determines the identity of the image and the face on which it is located. The images pasted on the cube faces were drawn from three: MNIST handwriting digit, QuickDraw, and RGB MNIST handwriting digit datasets. The attentional input of 3 concentric cropped windows resembling the high-resolution central fovea and low-resolution periphery of the retina, flows through a Classifier Network and a Camera Motion Network. The Classifier Network classifies the current view into one of the classes or clutter. The Camera Motion Network predicts the camera’s next position on the orbit (varying the azimuthal angle or ‘*θ*’). Here the camera performs one of three actions: move right, move left, or don’t move. The Camera-Position Network adds the camera’s current *θ* information into the higher features level of the Classifier Network and the Camera Motion Network. The Camera Motion Network is trained using Q-learning where the reward is 1 if the classifier network gives the correct classification, otherwise 0. Total loss is computed by adding the mean square loss of temporal difference and cross entropy loss. Then the total loss is backpropagated using Adam optimizer. Results on two grayscale image datasets and one RGB image dataset show that the proposed model is successfully able to discover the desired search pattern to find the target face on the cube, and also classify the target face accurately.

## I. Introduction

Human visual system (HVS) processes a restricted field of view of about 150 degrees in the horizontal line and 210 degrees in the vertical line [1]. However, the eye orientates itself in such a manner that the image of the region of interest falls inside the central part of the retina or fovea to obtain precise information from that part of the visual field. Information from the fovea in high resolution and periphery in low resolution is passed through the visual hierarchy, and the features related to the form, color, and motion are analyzed by respective visual cortical areas. Due to this anatomical constraint, the eye does not process the entire scene at once: the eye makes darting movements called saccades and attends the salient parts of the scene sequentially and integrates the pieces of the image to get a more comprehensive understanding of the scene.

Visual attention is a popular topic in both computer vision and visual neuroscience. A large number of computational models of visual attention, proposed in the past couple of decades: may be divided into two categories: Bottom-up approaches [2], [3], and top-down approaches [4], [5], [6]. The models are basically developed to predict the saliency map where a brighter pixel has higher probability of receiving human attention and vice versa. Bottom-up attention is considered to be stimulus driven whereas top-down attention is considered to be task driven which receives human attention based on the explicit understanding of the image content. Prior attempts in the field of top-down attention mechanisms [4], [5], [6] have mainly used nondeep networks such as a Bayesian approach [7] based on limited knowledge of visual attention. One of the popular study of the attention mechanism [8], Mnih et. al have developed an recurrent attention model (RAM) which takes a glimpse of the attention window as input and uses the internal state of the network to find the next location to focus on in a non-static environment. Their proposed network processes multiple glimpses of windows to attend a part of the image at different levels of resolutions. Training of their model is done by using the reinforcement learning approach for classification of MNIST dataset for modeling task-driven visual attention. Design of their network is based on fully connected layers, which leads to a rapid increase in computational cost with image size, and therefore the network is perhaps not feasible for more complex real world tasks such as search in a 3D environment.

There is an extensive number of research studies that demonstrate the application of attentional search methods to solve real world problems in 2D space such as image cropping [9], object recognition [10], object segmentation [11], [12], [13], and video understanding [14], [15], [16], [17], [18]. But the use of visual attention in a 3D environment is still relatively under-explored. Earliest work in 3D target search is the Shape Nets [19] where the objective was to voxelize the target and use deep belief networks for training and prediction. Minut and Mahadevan [20] use Q learning to identify the next movement of the camera (action) out of the eight possible actions in order to focus on the object of interest. At a lower level this approach uses histogram back projection color maps and symmetry map to identify the objects. Unlike reinforcement learning based approaches, the model proposed by Kanezaki et. al, named RotationNet [21], focuses on convolutional neural networks (CNNs) based approaches where each view of the object is taken into consideration for learning. The model predicts the class and the pose (orientation) of the object of consideration. This was an improvement over the previous CNN based networks which failed to predict the pose. The model yielded an accuracy of 94% on Modelnet40 dataset [19] consisting of 40 categories including chair, airplane etc. Multiview CNN [22] was one of the earliest attempts in 3D object recognition that acts as a precursor of the RotationNet. In the model known as the SaccadeNet [23], developed by Lan et. al, a model closest in approach to ours, four module classifiers are used to recognize objects. These modules are - center attentive module, the corner attentive module, the attention transitive module, and the aggregation attentive module. Each module works on identifying the main key points of the object of interest, perhaps the center, corners, attend object centers and bounding boxes. This technique works similar to the proposed saccade approach inspired by human visual search. The drawback is that it works mainly on 2D inputs. While performing a target search in a 3D environment, the model needs to predict the next location of the camera and identify the object that the camera is looking for. To perform such search tasks in 3D space, time is one of the constraints which depends on the network design and input. We propose an Attentional Search Model in 3D space (ASM-3D) that takes the attentional glimpse instead of the entire image. The design of the model contains convolutional layers instead of fully connected layers to extract features and contains Elman and Jordan recurrence layers as well as JK-flip-flop recurrence layer [24] instead of LSTM to integrate the temporal attention history in the network. To generate the attentional glimpse, a set of concentric attention windows is used by taking the inspiration from [28], [25], [26].

The proposed model has the following brain-inspired features: 1) it has separate channels for image classification and camera movement, analogous to the “What pathway” and “Where pathway” in HVS; 2) it incorporates three types of recurrence connections: a) Local recurrence connection of Elman type [27], b) Global recurrence connection of Jordan type [28], c) Flip-flop neurons [29] that are capable of storing information for a long time. In this study, we show that the ASM-3D is effectively able to learn task-specific strategies and identify the targets. Our simulation results successfully shows that an attention-based network can be an efficient approach in dealing with target searching tasks in a 3D environment, which is demonstrated by using 3D Cluttered MNIST Cube dataset, 3D Cluttered QuickDraw Cube dataset, and 3D Cluttered RGB MNIST Cube dataset.

## II. The Proposed Approach

### A. Environment Overview

The virtual environment used in this study is created using openGL [30] (Fig.2). The environment contains a cube placed at the origin of a spherical coordinate system and a camera placed on a circular orbit around the cube. On this orbit of radius ‘r’, the camera revolves around the cube, always looking inwards towards the center of the cube (Fig.3b). As the camera moves on the orbit, it processes the views of the cube it captures and searches for the face that has a target pattern displayed on it (Fig.4b). The possible movements of the camera on the orbit are: “move right” (*θ*^+^), “move left” (*θ*^-^), or “don’t move” (*θ*) (Fig.3).

**Fig. 1:**
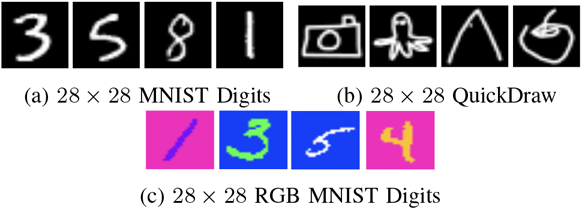
Sample of image datasets are shown here.

**Fig. 2:**
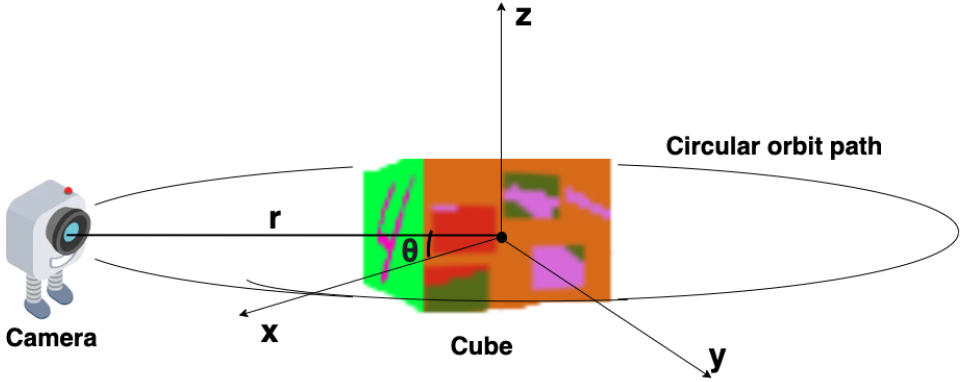
Simulated environment: ‘*r*’ is the radius of the circular orbit around the cube or the line connecting the camera position and origin of the spherical coordinate system. Polar angle ‘*ϕ*’ is assumed to be fixed at 0, and azimuth angle ‘*θ*’ is varying.

**Fig. 3:**
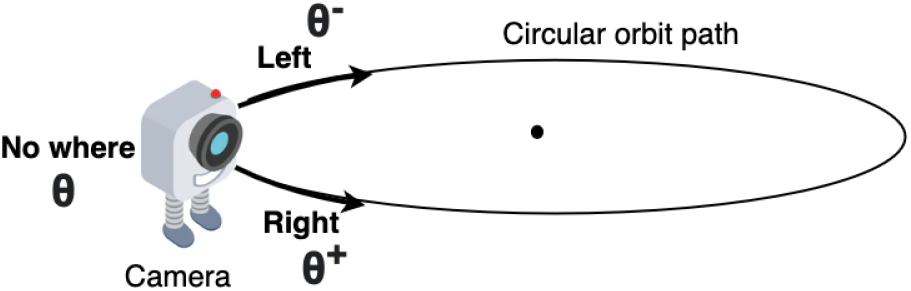
Direction of all 3 movements of the camera on the orbital path supposed to be predicted by the model

**Fig. 4:**
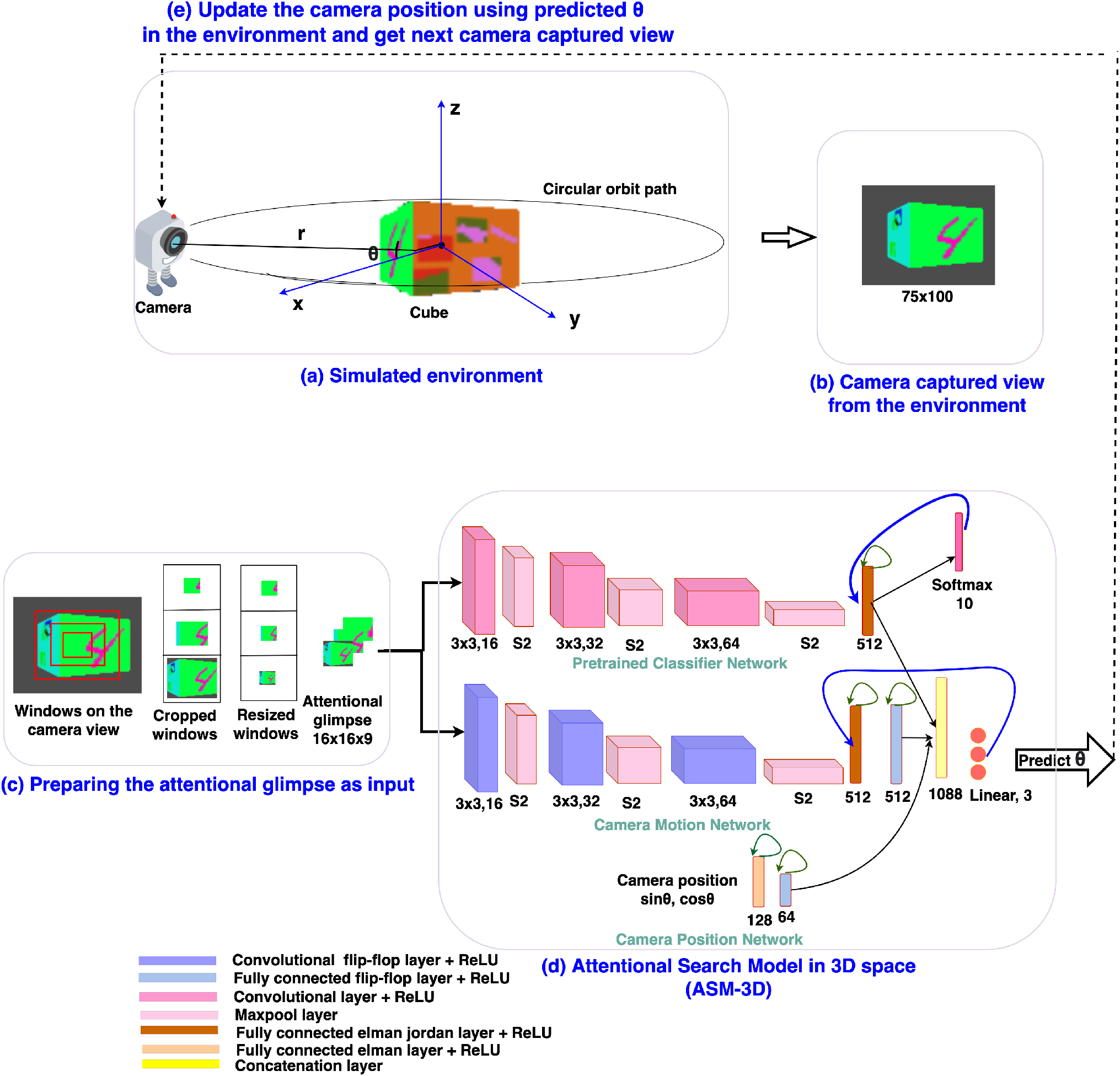
The design of the ASM-3D: (a) Simulated environment with the 3D Cluttered RGB MNIST cube and the camera, (b) Camera captures the image from the environment, (c) the attentional glimpse generated from the camera captured view, (d) the ASM-3D predicts the class of the target and position of the camera or ‘*θ*’, and (e) Update the camera position using the predicted ‘*θ*’.

### B. Architecture Overview

The architecture design of the proposed attentional search model in 3D space (ASM-3D) is depicted in Fig.4d. The model takes two inputs: i) the *attentional glimpse* which consists of the contents at different resolutions and sizes of the attended region, where multiple concentric attention windows are applied to the center location of the camera view, and ii) the *camera-position* in the form of a point on the unit circle at an angle *θ* or the azimuth angle of the camera movement in its circular orbit. The model predicts two outputs at each timestep: i) the next location of the camera on the orbit, and ii) the class of the object seen in the camera view. The model consists of three parallel pipelines (Fig.4d): i) the upper pipeline processes the class information of the object seen in the view, called the Classifier Network, ii) the middle pipeline processes the location of the target object over the cube, called the Camera Motion Network, and iii) the lower pipeline, which incorporates the camera position into the high level features of the Classifier Network and the Camera Motion Network, is called the Camera-Position Network. Outputs of all the three pipelines are concatenated in one flatten layer which connects with a fully connected layer and the output of the fully connected layer passes through one linear output layer and one softmax output layer in parallel. Linear output layer computes the Q-values corresponding to the movements in all three directions that can be taken by the camera and softmax output layer computes the class probabilities of the object present inside the attentional glimpse. Deep Q-learning algorithm is applied to train the model and learn the optimal policy for camera control. As the model takes the sequential input, the network requires memory to store the past information of the following details: i) the extracted features of the attentional glimpse, ii) its corresponding location on the cube, and iii) the camera position. For storing this input history, the model uses 3 recurrent neural features: the flip-flop neuron layer [29], Elman and Jordan recurrence layers.

### C. The ASM-3D

The proposed attention model is a deep neural network, which has three pipelines: Classifier Network, Camera Motion Network, and Camera-Position Network. The classifier network consists of three convolutional layers (Convs), three maxpool layers, and one fully connected (FC) Elman Jordan recurrence layer (FCEJ). The camera motion network consists of three convolutional flip-flop layers (ConvJKFF), three maxpool layers, one FCEJ layer, and one FC flipflop layer (FCJKFF). The camera-position network consists of one FCEJ layer, and one FCJKFF layer. This network encodes the revolving direction of the camera. The aforementioned layers are discussed in greater detail in the following paragraphs.

Convolutional layers (Convs) are used to extract feature by sharing the weights across different spatial locations. Input and output to the Conv layer are 3D tensors, called feature maps. The output feature map is calculated by convolving the input feature map with 3D linear filters. Then a bias term is added up into the convolved output. If **X**^*l*–1^ is the input feature map of *l^th^* Conv layer and **W**^*l*^ and **b**^*l*^ are filter weights and bias terms respectively, then the output feature map **X**^*l*^ of *l^th^* layer is calculated via equation-1:

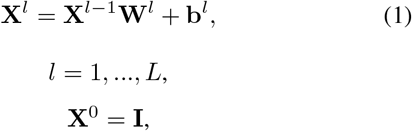

In the above equation, L is the total number of layers, **X**^0^ is the input image **I** to the first Conv layer. The output feature maps from each Conv layer are passed through a non-linear ReLU activation function [31] (equation-2).

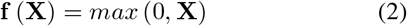

The output feature maps from the activation function, get normalized using local response normalization (LRN) [32]. LRN normalizes the feature maps within the channels that also implement lateral inhibition (equation-3).

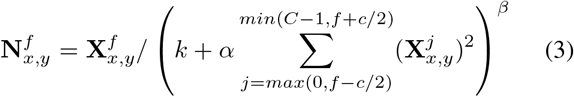

where **X**(*x, y*) and **N**(*x, y*) are the pixel values at (*x, y*) position before and after normalization respectively, *f* denotes the filter. *C* stands for the total number of channels. The constants *k, α, β*, and *c* are hyper-parameters. *k* is used to avoid “division by zero”, *α* is a normalization constant, while *β* is used as a contrasting constant. The constant *c* is used to define the length of the neighborhood, that is the number of consecutive pixel values need to be considered while calculating the normalization. (*k, α, β, c*) = (0, 1, 1, *C*) case is considered as the standard normalization. Normalized features from the Conv layer are passed through the maxpool layer [33]. Several convolutional layers and pooling layers are assembled alternately across depth in the first three Conv or ConvJKFF layers in both classifier and camera motion networks (Fig.4d).

To implement the Elman recurrence layer [27], the output vector of the FC layer at time ‘*t* – 1’ is stored in a context layer and the content of the context layer is fed back to the same FC layer at time ‘*t*’, named as FC Elman recurrence layer which is a short range storage connection. The Elman recurrence layer has been implemented only in the first FC layer of all three networks. Similarly, to implement the Jordan recurrence layer [28], the output vector of the last FC layer at time step ‘*t* – 1’ is stored in a context layer and this context layer is fed back to the first FC layer at time step ‘*t*’ in their corresponding pipeline, named as FC Jordan recurrence layer which is a long-range storage connection. In this way the first FC layer in Classifier and Camera Motion Networks has both Elman and Jordan recurrences; so we call this layer a FCEJ layer. The computation of FCEJ is shown in the following equation-4

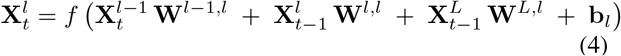

In equation-4, 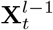 is the output of the *l* – *1^th^* layer at time ‘*t*’ and going as input to the *l^th^* layer at time ‘*t*’ (FC layer). 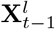 is the output of the *l^th^* layer at time ‘*t* − 1’ and going as input to the same *l^th^* layer at time ‘*t*’ (Elman recurrence layer). 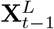 is the output of the *L^th^* layer at time ‘*t* – 1’ and going as input to the *l^th^* layer at time ‘*t*’ (Jordan recurrence layer). **W**′*s* and *b* are the corresponding weights and bias respectively. *f* is the ReLU activation function.

Memory of the past information in the layers of the proposed network is stored using a third mechanism – the flip-flop neurons [29]. A flip-flop is a digital electronic circuit to store state information. There are 4 types of digital implementations of flip-flops: D flip-flops, Toggle flipflops, SR flip-flops, and JK flip-flops [34]. In the proposed network, JK flip-flop neurons have been used in place of LSTM neurons because of the performance advantage shown in [24], and [29]. In both of these papers, the experiment conducted on the sequential data shows that flip-flop neurons outperform the LSTM neurons, using only half the number of training parameters in comparison to LSTM. Likewise, to get the advantage of less parameters and better performance, in the current study we used the JK flip flop neuron. The JK flip-flop neuron uses 2 gating variables with “**J** and **K**” nodes whereas LSTM uses 4 gating variables. In this paper, the term flip-flop will be used to refer to JK-flip-flop. Furthermore, the flip-flop neurons are considered similar to the UP/DOWN neurons found in the prefrontal cortex (PFC), responsible for working memory [35].

In the proposed model, the flip-flop layer is designed in two ways: flip-flop neurons in convolutional layer (named as “convolutional flip-flop layer” or ConvJKFF), and flip-flop neurons in the FC layer (named as “fully connected flip-flop layer” or FCJKFF). Training rules of these flip-flop neurons in the network were also developed. The two gate outputs ‘**J**’ and ‘**K**’, the hidden state of the JK flip-flops, and the final flip-flops output are computed by using eqns (5, 6, and 7 respectively) below.

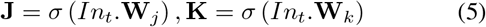

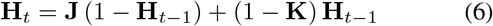

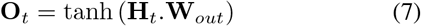

In eqns (5 & 7), ‘.’ stands for the matrix multiplication in case of FCJKFF layer whereas the convolve operation in case of ConvJKFF layer. *In_t_* = (**X**_*t*_; **H**_*t*_) is the input to the flip-flop layer, where **X**_*t*_ is the output from previous layer and **H**_*t*_ is the hidden state at time ‘*t*’, which initialize with ones at time 0. **J** and **K** are the gate variables, which has weight parameters **W**_*j*_ and **W**_*k*_ respectively. **O**_*t*_ is the output of the flip-flop layer at time ‘*t*’. To train the flip-flop neurons, the partial derivatives w.r.t **J** and **K** were used to backpropagate the corresponding **J** and **K** nodes (equation-8).

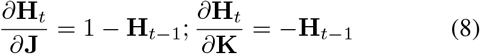

### D. Implementation Detail

#### 1) Camera Motion Network

The camera motion network takes the attentional glimpse of size *h* × *w* × *a* as input, where ‘*h*’ is the height, ‘*w*’ is the width, and ‘*a*’ is the number of the cropped attention windows. Here, the number of attention windows is chosen to be 3 (i.e, *a* = 3). The size of one attention window is twice the previous attention window’s size. Similar multi-scale concentric attention windows were used in other models [8], [36], [37]. All of the attention windows, except the smallest one, get resized to the size of the smallest attention window. For example, to generate the attentional glimpse where *h* = 16, *w* = 16, and *a* = 3 from location *y* = 35 and *x* = 50 in the given image of size 75 × 100, the first, second, and third attention windows are cropped out of size 16 × 16, 32 × 32, and 50 × 50 from pixel location (*y, x*) = (27 to 43, 42 to 58), (*y, x*) = (19 to 51, 34 to 66), and (*y, x*) = (10 to 60, 25 to 75) respectively. The second and third cropped attention windows are resized into the size of the first cropped attention window, which is 16 × 16. After resizing, all of the three attention windows are stacked together which finally becomes an attentional glimpse of size 16 × 16 × 3. This type of attentional glimpse having a size of *h* × *w* × *a* shown in Fig.4c is passed to the first ConvJKFF layer of 16 kernels, each of size 3 × 3, of the classifier network (shown in the top pipeline of the ASM-3D in Fig.4d). The spatial dimension of the features generated from the first ConvJKFF layer is *h* × *w* × 16, which are normalized using LRN, and passed into ReLU activation function. Output from ReLU activation function is passed to the maxpool layer with a window of size 2 × 2 and stride by 2, which translates the feature’s spatial dimensions into *h*/2 ×*w*/2 × 16. The translated feature maps are passed as input to the second ConvJKFF layer of 32 kernels, each of size 3 × 3, to extract the higher level features of size *h*/2× *w*/2 × 32. Then, similar to the previous layer, features generated from the second ConvJKFF layer are passed through the LRN layer, ReLU activation function, and maxpool layer with a window of size 2 × 2 and stride 2. After passing into the maxpool layer, feature maps of size *h*/4 × *w*/4 × 32 are generated, which further goes to the flattened layer to reshape the 3D features into 1D vectors. The flattened vectors are passed through one FCEJ layer of 512 neurons, which is followed by one FCJKFF layer of 512 neurons. Output from the FCJKFF layer of the camera motion network is concatenated with the output vectors of the last layer of the other two channels.

#### 2) Classifier Network

The classifier network gets the same attentional glimpse as input which has been passed to the camera motion network. This network predicts the class of the object present in the attentional glimpse. The object present in the attentional glimpse may belong to one of the ‘*n*+1’ classes, where ‘*n*’ classes are the object or target class and one is the nontarget or clutter class. The network consists of 3 Conv layers followed by one FCEJ layer. The first Conv layer of 16 kernels of size 3×3 generates the feature maps of spatial dimension *h* × *w* × 16. Generated features are passed through the LRN layer and ReLU activation function. After this, the maxpool layer with a window of size 2 × 2 and stride by 2 has been applied to the output of ReLU activation function, which gives the feature maps of spatial dimension *h*/2 × *w*/2 × 16. Then, the feature maps are passed through a second Conv layer of 32 kernels, each of size 3 × 3, LRN layer, ReLU activation function, and maxpool layer with a window of size 2 × 2 and stride by 2. Feature maps of spatial dimension *h*/4 × *w*/4 × 32 are passed through a third Conv layer of 64 kernels each of size 3 × 3 with ReLU activation function, which further generates the feature maps of size *h*/4 × *w*/4 × 64. Then the flattened layer reshapes the 3D tensor of feature maps into vectors and these vectors are input to the FCEJ layer of 512 neurons. Output of the FCEJ layer gets concatenated with the output vectors of the last layer of the camera motion network and the camera position network.

#### 3) Camera Position Network

Camera’s position in the environment is inferred from the spherical coordinates, where the camera is assumed, as described before, on a circular object centered on the origin, and the center of the cube is located at the origin. The camera’s position is defined from using three variables: (‘*r*’, ‘*θ*’, ‘*ϕ*’), where ‘*r* = *R*’ is the radius of the sphere or line connecting the camera point and the origin of the spherical coordinate system, *θ* is the azimuth angle and ‘*ϕ*‘ is polar angle of the spherical coordinate system. In the current simulated environment, the camera moves only in one degree of freedom, that is ‘*θ*’. Therefore, ‘*r* = *R*’ and ‘*ϕ* = 0’ are considered to be constant. Since only *θ* varies as the camera moves on a circular orbital path around the origin of the spherical coordinate system or the cube. In the camera position network, sinusoidal waves of *θ* are passed as input to the first FC Elman (FCE) layer having 128 neurons followed by one FCJKFF of 64 neurons. Output from the FCJKFF layer is concatenated to the output vectors of the last layer of the classifier network and the camera motion network.

Outputs from three pipelines are concatenated in one common flattened layer, which further connects with 2 output layers in parallel. One output layer with linear activation function is responsible to predict one direction out of the 3 considered directions in which the camera will move on the orbit to look and locate the target face present in the given cube. The other output layer with softmax activation function is responsible to predict the class of the object seen on the view of the camera.

#### 4) Training and Testing

Tensorflow framework is used to implement the proposed attention model. Xavier initialization [38] (equation-9) with random normal distribution is used to initialize the weights for each layer of the three networks. The Xavier initialization is able to avoid the exploding or vanishing gradients [39] problem by fixing the variance of the activations across each layer as the same.

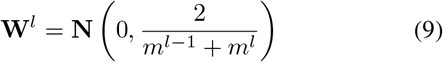

where, **N** stands for the normal distribution. *m*^*l*–1^ and *m^l^* is the number of neurons in the previous layer and current layer respectively. **W**^*l*^ denotes the weights at *l^th^* layer with Xavier initialization.

Before training the model, the classifier network is pretrained on the camera captured views. To pretrain the classifier network, we collect views of the simulated environment by explicitly revolving the camera from −180 to +180 degree where 0 degree is assumed to be exactly at the front of the face containing the target object. Advancing in steps of 9 degrees over the range of −180 to +180 degree, a total 40 views are collected for each cube in the dataset. Views between −45 to +45 range are labeled as one of the ‘*n* classes’ and views between +46 to +180 and −46 to −180 range are labeled as ‘background class’. Therefore, total number of classes present in the dataset is *n*+1. To make the views data uniform, the same number of views of the background class are chosen randomly as the number of views of the other class. The classifier network is pretrained on such views of targets and background or nontarget class so that the classifier network in the model has the knowledge to differentiate between the target class and nontarget class. We assume that the camera’s focus is always fixed on the center of the view. Therefore, we create a glimpse of three concentric windows from the center location of the camera view. Detailed architecture of the pretraining classifier network is shown in Fig.5. The classifier network without recurrent layers in the ASM-3D is pretrained on the glimpse of the camera views. Total loss of the model is calculated in two parts: one is classification loss, calculated using the cross-entropy loss function [40] and the other one is prediction loss, calculated using mean square error of temporal difference [41]. Equations of the both loss functions are shown in equation-10 and 13.

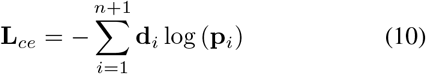

**Fig. 5.**
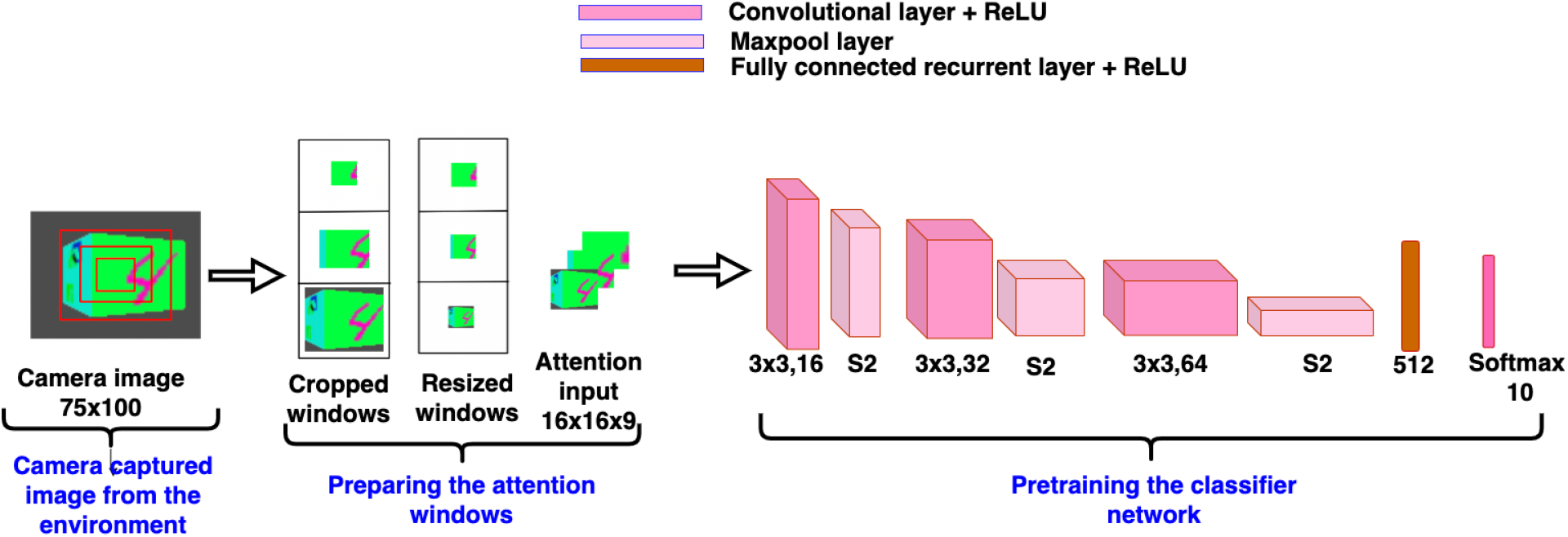
Pretrained classifier network

In equation-10, **d**_*i*_ denotes the desired class probability and **p**_*i*_ denotes the predicted class probability of *i^th^* class. ‘*n*+1’ is the total number of classes that are present in the dataset. Here, the camera is assumed as an agent and the agent learns a defined policy of the reward function (equation-11) [42]. When the agent is in the current state, Q-values of all three actions are predicted by passing the information of the current state (like the attention input and the *θ* value of the camera) into the deep neural network. Based on the predicted Q-value of all the actions in the current state, the agent makes an action decision using a softmax action selection policy [43]. In this policy, the predicted Q-values are passed through a softmax activation function to produce the action probabilities. The action with the highest probability is selected and performed by the agent in the current state of the environment. After performing the action, the current state is updated to the next state and then the agent receives a reward, either ‘1’ or ‘0’ depending on the reward policy shown in equation-11.

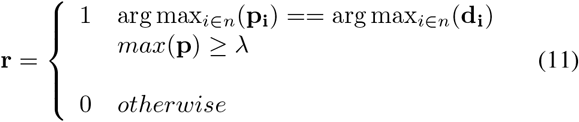

Apart from the softmax action selection policy, we used a race model [44] which ensures that the selected action is correct. Race models have been applied in many behavioral, perceptual, and oculomotor decisions and such decisions are based on trial-to-trial modifications in a race among all the responses [45]. Race model works based on two neurophysiological evidences to show the relatedness. Firstly, if monkeys are trained to make their decision on coherently moving direction of dots, accumulating neuronal activity is formed that mirrors the decision even when there is no coherent motion. Here, both choices are equally rewarded [46]. Secondly, the decision threshold is considered constant for a selected action, regardless of its being a specifically cued action [47]. We have taken the motivation to apply the race model based on the second evidence. The action predicted by the network is the action which crosses the threshold λ, first and if the action predicted is correct, the agent gets reward ‘1’; it otherwise gets reward ‘0’.

The Q-values of the actions in the next step are estimated by passing the next state information into the target network, where the target network is the separate copy of the networks of the model. Target Q-value is calculated by adding the current state reward and maximum of the next state Q-values multiplied with a discount factor *γ*. Discount factor defines how much the current state Q-value depends on the future reward. Now the temporal difference (TD) is calculated by calculating the difference between the target Q-value and the predicted Q-value (equation-12).

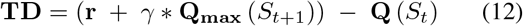

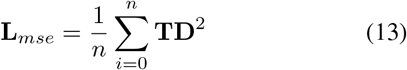

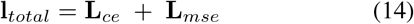

where, **r** is the reward which the agent gets while going from the state **S**_*t*_ to the state **S**_*t*+1_. **Q**(*S*_*t*+1_) and **Q**(*S_t_*) is the Q-value of the state **S**_*t*+1_ and **S**_*t*_ respectively. *γ* is the discount factor. Then these two losses, the cross-entropy loss of the classifier network (equation-10), **L**_*ce*_ and the mean square error of temporal difference of the camera motion network (equation-13), **L**_*mse*_, are added up to get the total loss (equation-14). The total loss is back propagated into the network [48]. The network parameters are updated by using the mini-batch Adam optimizer [49]. L2 regularization [50] has been used to avoid the overfitting problem of the network. During inference, the camera starts from a random location and moves towards the target face of the cube. Once it finds the target face, the camera continues to fixate around that face. The model achieves a processing speed of 0.0187 seconds per input image on a workstation with a NVIDIA GeForce GTX 1080Ti 11GB GPU and 32GB RAM.

## III. Simulation Results

We evaluate our model on “painted cube” data, where each cube has a target object on one vertical face and nontarget objects on the other three vertical faces. The model is supposed to move the camera around the cube on a circular orbit and search the target object image present on one of the four vertical faces of the cube. For target object image, we have used image datasets. Totally three 3D Cluttered Cube datasets have been considered in the experiment. Each of the cube datasets has been generated using their related image data. Grayscale MNIST digit image dataset, QuickDraw image dataset, and RGB MNIST digit image dataset were used to generate cube datasets like 3D Cluttered Grayscale MNIST Cube dataset, 3D Cluttered QuickDraw Cube dataset, and 3D Cluttered RGB MNIST Cube dataset respectively. First two of these are cube datasets with grayscale images and the last one is a cube dataset with RGB images. Based on the grayscale and RGB cube datasets, we have designed the experiments in two parts: one part of the experiment shows the target search capability of the proposed model on the cubes which has all 4 vertical faces of grayscale images (called grayscale cubes) and the other part of the experiment shows the target search capability of the model on the cubes which has all vertical faces of RGB images (called RGB cubes).

### A. Searching on Grayscale Cubes

In the first part of the experiment, we evaluated our model on 2 datasets of Grayscale cubes. For that we used 2 different datasets of grayscale images: MNIST handwritten digits [51] and QuickDraw [52]. Both datasets with 10 different classes contain 48, 000 examples in the training set, 12, 000 examples in the validation set, and 10, 000 examples in the testing set. We have generated a 3D Cluttered MNIST Cube dataset using MNIST dataset. To generate such a cube dataset, each of the cubes were created with a 28 × 28 MNIST digit image (target) on one vertical face and 28 × 28 random clutter image (non-target) on the other three vertical faces. In this experiment, the bottom and top faces of the cube are not considered for searching. Similarly, a 3D Cluttered QuickDraw Cube dataset was generated using QuickDraw image dataset.

Once the cube datasets are generated, we place the cube in the environment in such a way that the center of the cube is at the origin of the spherical coordinate system. Then the camera is placed at a random value of azimuth angle ‘*θ*’ at initial time (*t* = 0). The polar angle ‘*ϕ*’, and radius ‘*r*’ are set to 0, and 2.5 respectively. The camera placed at (*r, θ, ϕ*) captures the view of size 75 × 100. Then a glimpse is extracted from the center location of the captured view. To extract the glimpse, three concentric windows of size 16 × 16, 32 × 32, and 50 × 50 are cropped out from the center of the view. After cropping out, windows of size 32 × 32 and 50 × 50 are resized into the size of 16 × 16. Then resized windows with the smallest window of size 16 × 16 are arranged together across depth to generate attentional glimpse of dimension 16 × 16 × 3. Since the image size in the QuickDraw image dataset is same as the image size in the MNIST dataset, the same dimensions of the camera view and attentional glimpse were considered in case of the 3D Cluttered QuickDraw Cube dataset.

The proposed model takes the attentional glimpse of size 16 × 16 × 3 from the center location of the image view of the camera of size 75 × 100. Achieved accuracy on both grayscale datasets are listed in Table-I. The results of the camera’s movement predicted by our model in the testing set are shown in Figs.6, and 7. In this figure, images of the camera view of dimension 75×100 are shown in one row and their corresponding plots for predicted class probabilities for that view (dotted dashed-blue curve) and ground truth target class probabilities (green curve) are shown in the row just below. At the bottom of the plots, timestep and ground truth target class labels are denoted by using variables ‘*t*’ and ‘*c*’ respectively. In the row of images of the camera view, three concentric red windows depict the glimpse.

**TABLE I:** Accuracy on testing set of all three datasets

| Dataset | Testing Accuracy |
| --- | --- |
| 3D Cluttered Grayscale MNIST Cube dataset | 95.6% |
| 3D Cluttered Grayscale QuickDraw Cube dataset | 83% |
| 3D Cluttered RGB MNIST Cube dataset | 91.5% |

**Fig. 6:**
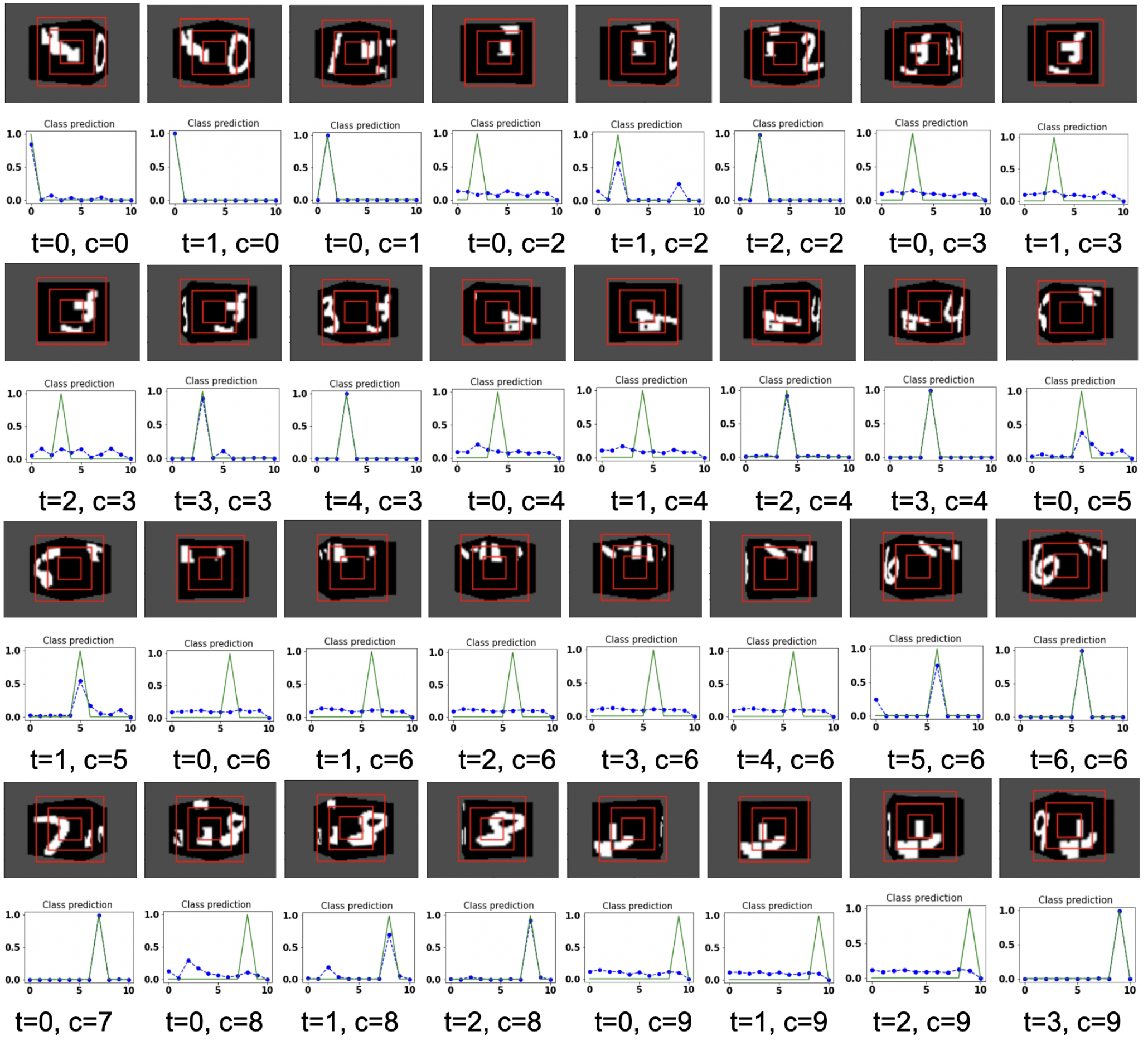
Illustrates the camera movements around the cube to search the target face in the view of size 75 × 100, predicted by our model in 3D Cluttered Grayscale MNIST Cube dataset. For each class at time *t*, there is a movement (shown in the row of camera view images) and corresponding classification probabilities (shown in the row of plots). In the row of camera view images, the three concentric red windows depict the glimpse at the center of the view image. In the plot corresponding with the above view image, green curve is the desired class probabilities and dotted dashed-blue curve is the predicted class probabilities at time *t*.

**Fig. 7:**
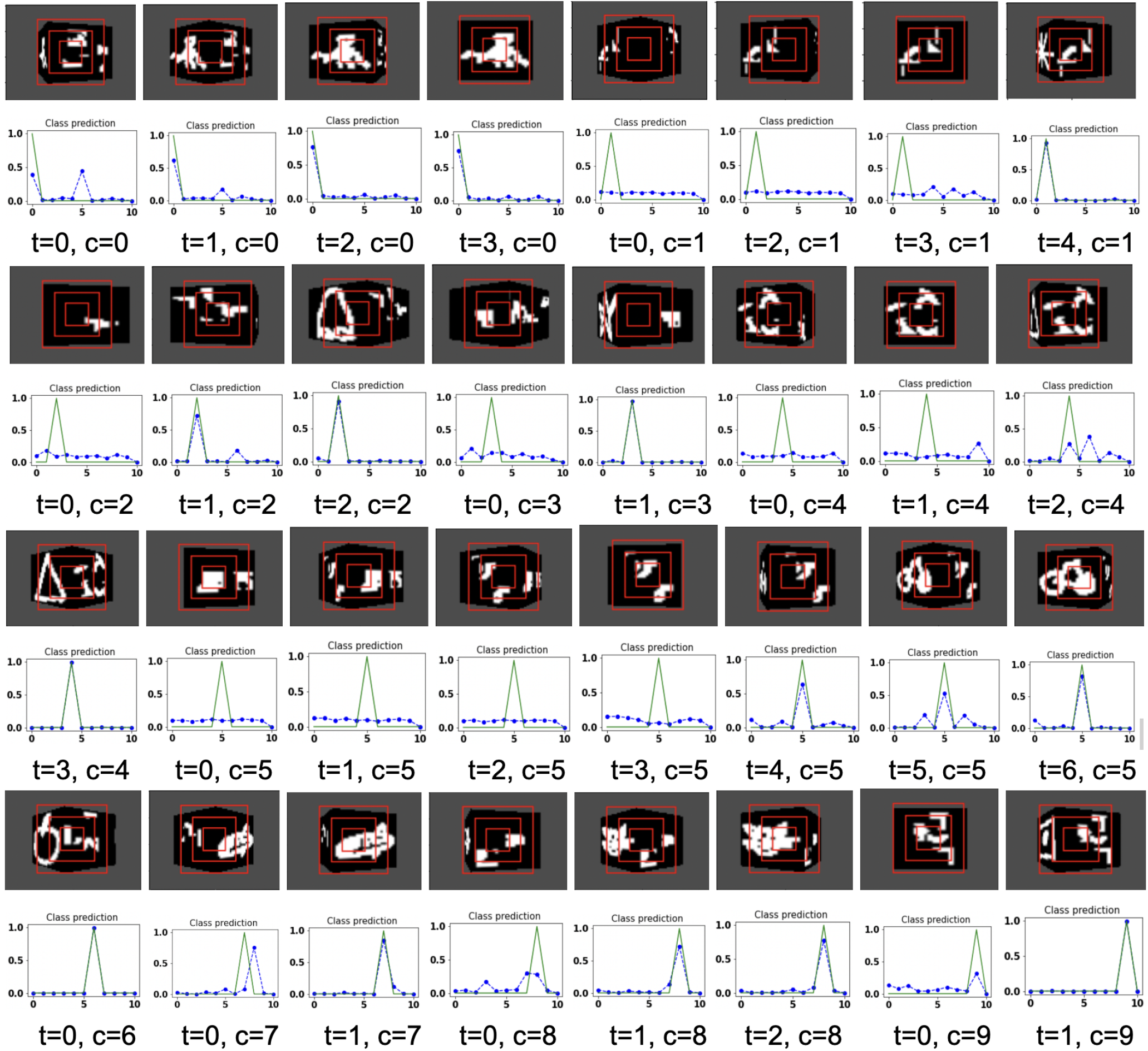
Illustrates the camera movements around the cube to search the target face in the view of size 75 × 100, predicted by our model in 3D Cluttered Grayscale QuickDraw Cube dataset. For each class at time *t*, there is a movement (shown in the row of camera view images) and corresponding classification probabilities (shown in the row of plots). In the row of camera view images, the three concentric red windows depict the glimpse at the center of the view image. In the plot corresponding with the above view image, green curve is the desired class probabilities and dotted dashed-blue curve is the predicted class probabilities at time *t*.

The model has the ability to move the camera in the position where the target face of the cube is visible from the camera. For example, in Fig.6, the class of digit 2 in the fourth image of the first row has the view of nontarget or clutter face at timestep *t* = 0 and its corresponding predicted class probabilities shown in the plot just below to that image is low for all classes. But at timestep *t* = 1 (*θ* is decided by the model), the camera has moved towards the right and has seen some part of the target face that has the digit 2. At the same time, the highest of the predicted class probabilities is for digit 2. The camera is again moved towards the right direction at timestep *t* = 2 and maximum value of predicted class probabilities is close to 1. The search for the target face ends when an adequate part of the digit 2 on the face is visible to the camera. Similarly for the other digits, the camera starts moving appropriately and searches for the target. The camera stops moving when an adequate part of the target is visible and the maximum value of the predicted class probabilities crosses a threshold of value 0.95. Threshold is set based on the feature complexity of the image datasets.

In the case of the 3D Cluttered QuickDraw Cube dataset, we can observe the same search behavior of the camera. For example, in Fig.7, class 5 (bicycle) in the second image of the seventh row has the camera view showing nontarget objects on the cube face at timestep *t* = 0 and its corresponding predicted class probabilities shown in the plot just below to that image is low for all classes. At the next timestep (*t* = 1), the camera has moved towards the left and the camera continues to move in the left direction 3 more times even though the target is not visible. At timestep *t* = 4, a very small part of the bicycle is visible and at this time the class probability for class 5 or bicycle becomes the highest. The camera stops moving once the maximum value of the predicted class probabilities crosses a threshold of value 0.85.

### B. Searching on RGB Cubes

In the second part of the experiment, we evaluated our model on RGB cubes to investigate that the model is able to search for the target object on the cube face even in the case of color images. To this end, we generated a cube dataset using RGB MNIST image dataset. Here, we first create the RGB MNIST digit image dataset by assigning different colors to the digits and the background of the images available in Grayscale MNIST digit image dataset [51]. The dataset with 10 different classes contains 48, 000 examples in the training set, 12, 000 examples in the validation set, and 10, 000 examples in the testing set. Once the image dataset is ready, we generate a 3D Cluttered RGB MNIST Cube dataset using RGB MNIST image dataset. To generate a 3D Cluttered RGB MNIST Cube dataset, each of the cubes is created with a 28 × 28 × 3 RGB MNIST image (target) on one vertical face and 28 × 28 × 3 random clutter image (non-target) on the other three vertical faces.

The model is evaluated by placing the RGB cube in the environment in the same way of grayscale cube datasets. The camera captures the view of size 75 × 100 of the 3D Cluttered RGB MNIST Cube. The camera extracts the glimpse from the center of the captured view. To extract the glimpse, three concentric windows of size 16 × 16, 32 × 32, and 50 × 50 are cropped out from the center of the view to generate attentional glimpse of size 16 × 16 × 9. The proposed model takes the attentional glimpse of size 16 × 16 × 9 from the center location of the image view of the camera of size 75 × 100 in case of 3D Cluttered RGB MNIST Cube dataset. The achieved accuracy on the RGB cube dataset is listed in Table-I. The results of the camera’s movement predicted by our attention model in the testing set are shown in Figs.8, and 9. Plots for accuracy, and reward vs. epoch is shown in Fig.10.

**Fig. 8:**
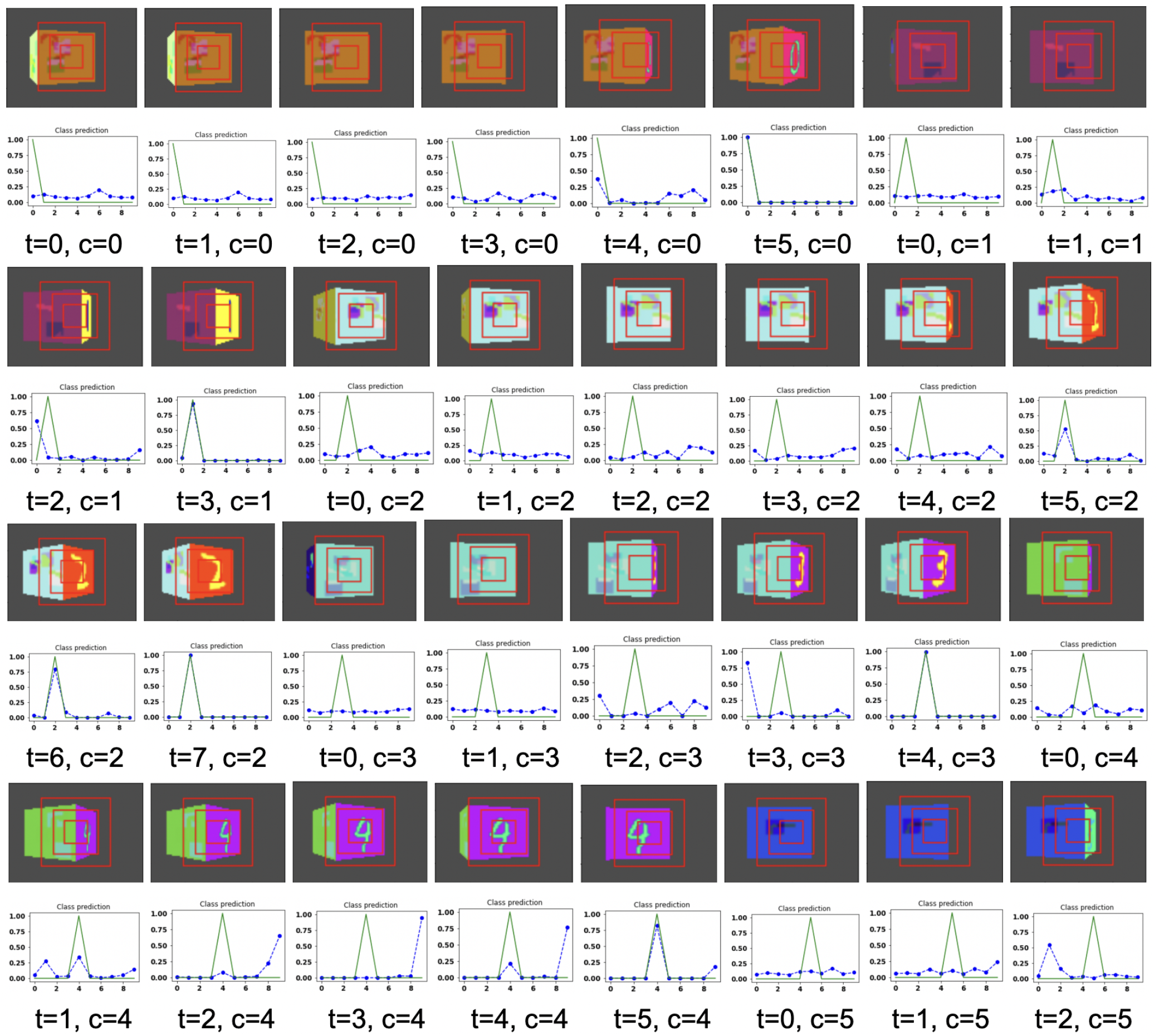
Illustrates the camera movements around the cube to search the target face in the view of size 75 × 100, predicted by our model in 3D Cluttered RGB MNIST Cube dataset. For each class at time *t*, there is a movement (shown in the row of camera view images) and corresponding classification probabilities (shown in the row of plots). In the row of camera view images, the three concentric red windows depict the glimpse at the center of the view image. In the plot corresponding with the above view image, green curve is the desired class probabilities and dotted dashed-blue curve is the predicted class probabilities at time *t*.

**Fig. 9:**
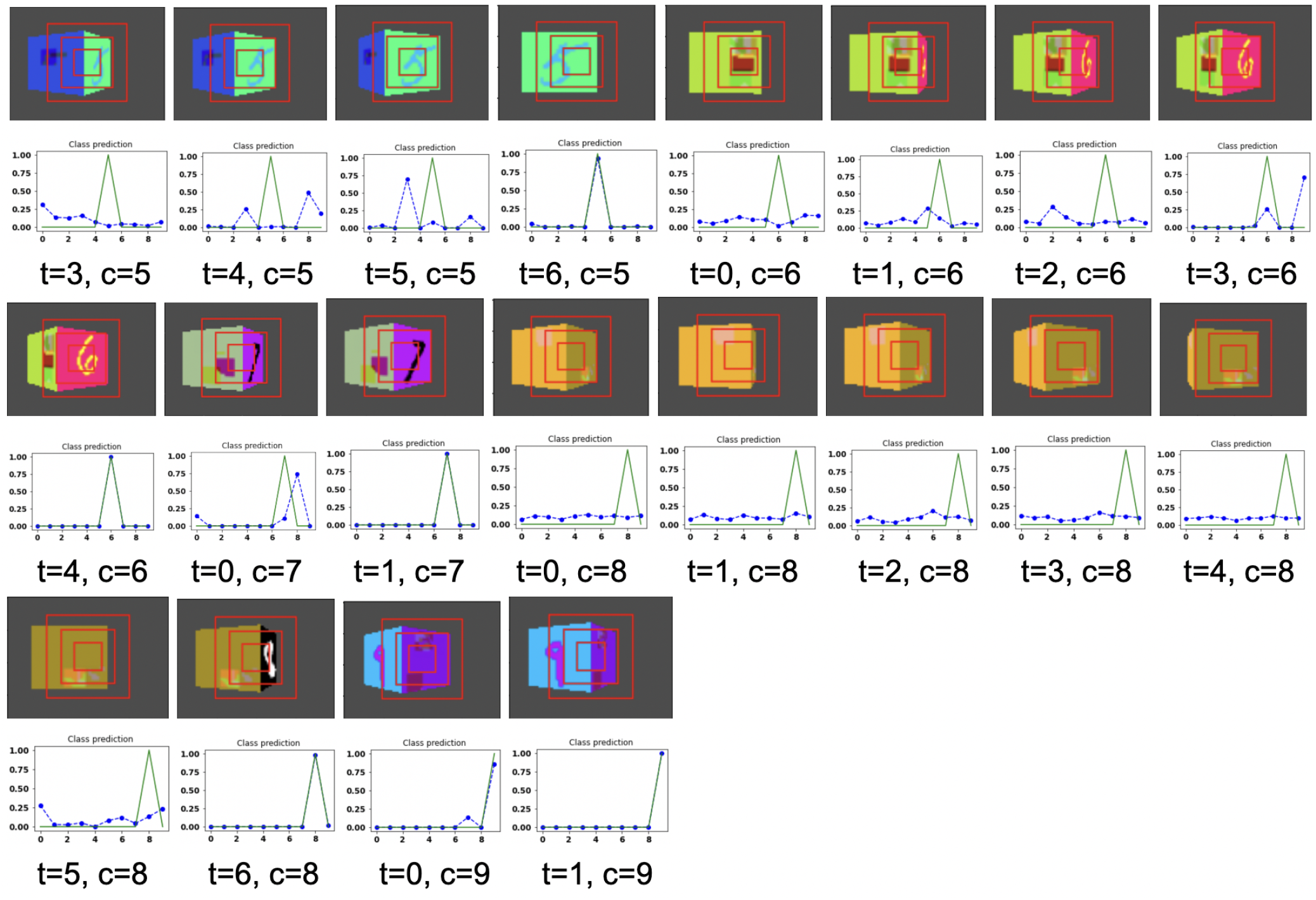
Illustrates the camera movements around the cube to search the target face in the view of size 75 × 100, predicted by our model in 3D Cluttered RGB MNIST Cube dataset. For each class at time *t*, there is a movement (shown in the row of camera view images) and corresponding classification probabilities (shown in the row of plots). In the row of camera view images, the three concentric red windows depict the glimpse at the center of the view image. In the plot corresponding with the above view image, green curve is the desired class probabilities and dotted dashed-blue curve is the predicted class probabilities at time *t*.

**Fig. 10:**
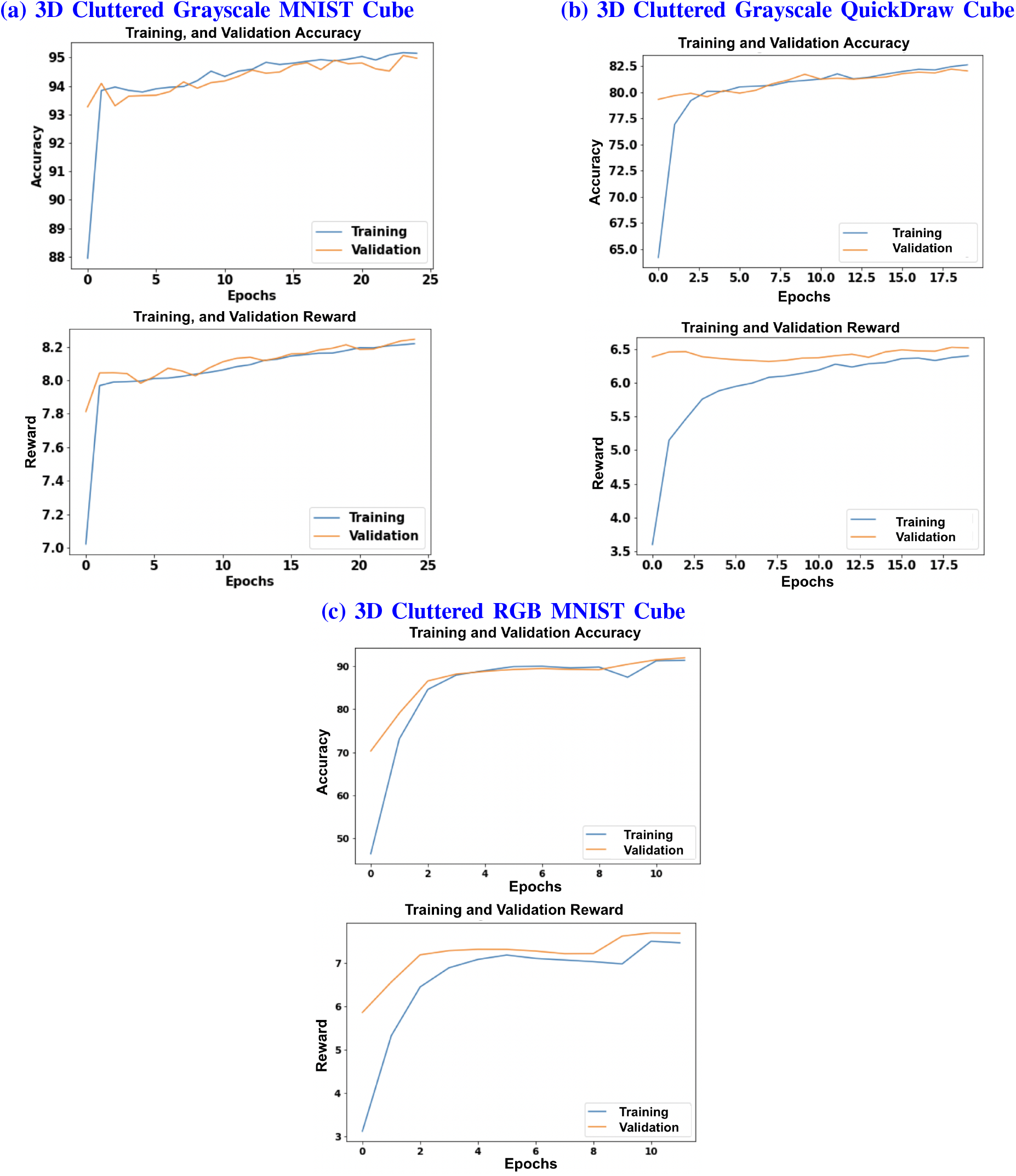
Column (a), (b), and (c) shows the plots of Accuracy (1^*st*^ and 3^*rd*^ row) and Reward (2^*nd*^ and 4^*th*^ row) vs Epochs of 3D Cluttered Grayscale MNIST Cube dataset, 3D Cluttered Grayscale QuickDraw Cube dataset, and 3D Cluttered RGB MNIST Cube dataset respectively.

The hyperparameters of the model are tuned and chosen as follows: 0.0001 learning rate, 0.43 discount factor, 0.85 lambda, and 0.1 regularization factor with the best performance in case of 3D Cluttered Grayscale MNIST Cube dataset. The model explores the actions with *∊* equal to 0.99 and the exploration gets reduced by a decay factor of 0.999 while training. The minimum value of *∊* is set with 0.1. The model is trained for 25 epochs and 50 timesteps, per cube in case of 3D Cluttered Grayscale MNIST Cube dataset. In the case of the 3D Cluttered QuickDraw Cube dataset, the model is trained for 20 epochs and 50 timesteps. During the inference, time-steps are varied depending upon the class probabilities. Prediction is considered to be done as soon as the maximum value of the class probabilities crosses a certain threshold (= 0.95). A slight variation in values of the hyperparameters is used for the 3D Cluttered RGB MNIST Cube dataset after tuning.

Jump length is the displacement from one location to the next location. The jump length of the camera from one location to the next location on the orbit is considered as a predefined parameter. The jump length of the camera is 12 in case of Grayscale 3D Cluttered MNIST Cube dataset, and 20 in case of 3D Cluttered QuickDraw Cube dataset and 3D Cluttered RGB MNIST Cube dataset.

## IV. DISCUSSION

To search for the entrance of a building, where there is neither a boundary wall, nor a clear path leading to the entrance, we usually move on the circular path around the building in either clockwise or anticlockwise direction until we find the entrance. While performing such a task, we also take care that the movement should not involve rapid alternation between the two directions, and must progress continuously in one direction. The best application of the current model can be in space. For example, geostationary satellites and spy satellites revolving around the earth in a circular orbit require a searching capability of one specific large area of the earth to collect bird’s eye view or to obtain the information about various weather, natural calamities, deforestation, and similar activities. From the results of camera movement shown in Figs.6, 7, 8 and 9, the proposed model is able to avoid alternative movements and is always able to follow the continuous movements to search the target face of the cube. There are three major components to consider the proposed model biologically inspired. First, the model takes the input of multiple concentric windows of different scales which resembles the differential spatial resolution of the central fovea and the peripheral regions of the retinal. Second, the model processes the view and its corresponding functions of the camera’s location, θ, which is analogous to determining the position using path integration and using it to navigate the world. The classifier and camera motion networks are analogous to the processing of visual information along the “what and where/how” pathways [53] respectively. Third, the model uses Elman, Jordan, JK-flip-flop recurrence layers as memory to store the history of the view and corresponding location, which resemble the feedback loops present among the visual cortical areas [54]. The output layers of the classifier and the camera motion network are used to attribute a specialized role to both of the networks for classification and searching tasks, by feeding the outputs back into the first fully connected Elman and Jordan layers in their corresponding channels. The output vector of the camera motion network (Q-values) which has information about the action to be taken by the camera is fed back into the fully connected Elman and Jordan layer and the output vectors of this layer passed through fully connected flip-flop layer and gets concatenated with the output of the last layer of the camera position network, this wide loop is responsible for storing the history of location and view.

## V. CONCLUSIONS

In the proposed model, we have shown how the “classifier” and “camera motion” networks coordinate with each other to perform the 3D visual search task. The ASM-3D successfully performed the classification task on a 3D environment on three datasets (Table-I). As shown in the results, movements generated by the model to search a target in the given cube always aim at the target face and take meaningful movements so that the camera looks at the target and classifies it correctly. Based on the results described herewith, we want to extend the model to more complicated full 3D searches in a 3D environment, like for example, searching for defects on the surface of a 3D structure. The model can then be applied to full scale object detection and recognition in 3D space.

## Acknowledgment

We acknowledge the support from Pavan Holla and Vigneswaran for the implementation of flip-flop neurons. We also acknowledge Sowmya Manojna to generate RGB MNIST dataset. Sweta Kumari acknowledges the financial support of Ministry of Human Resource Development for graduate assistantship.

